# CsMYB4a from *Camellia sinensis* Regulates the Auxin Signaling Pathway by Interacting with CsIAA4

**DOI:** 10.1101/2021.10.11.463959

**Authors:** Guo-Liang Ma, Ying-Ling Wu, Chang-Juan Jiang, Yi-Fan Chen, Da-Wei Xing, Yue Zhao, Ya-Jun Liu, Tao Xia, Li-Ping Gao

**Affiliations:** School of Life Science, Anhui Agricultural University, West 130 Changjiang Road, Hefei 230036 Anhui, China; State Key Laboratory of Tea Plant Biology and Utilization / Key Laboratory of Tea Biology and Tea Processing of Ministry of Agriculture / Anhui Provincial Laboratory of Tea Plant Biology and Utilization, Anhui Agricultural University, West 130 Changjiang Road, Hefei 230036 Anhui, China

## Abstract

Members of the R2R3-MYB4 subgroup are well-known negative regulatory transcription factors of phenylpropane and lignin pathways. In this study, we found that transgenic tobacco plants overexpressing a R2R3-MYB4 subgroup gene from *Camellia sinensis* (*CsMYB4a*) showed inhibited growth that was not regulated by phenylpropane and lignin pathways, and these plants exhibited altered sensitivity to synthetic auxin 1-naphthaleneacetic acid (α-NAA) treatment. An auxin/indole-3-acetic acid 4 (*AUX/IAA4*) gene from *Camellia sinensis* (*CsIAA4*) participating in the regulation of the auxin signal transduction pathway was screened from the yeast two-hybrid library with CsMYB4a as the bait protein, and tobacco plants overexpressing this gene showed a series of auxin-deficiency phenotypes, such as dwarfism, small leaves, reduced lateral roots, and a shorter primary root. *CsIAA4* transgenic tobacco plants were less sensitive to exogenous α-NAA than control plants, which was consistent with the findings for *CsMYB4a* transgenic tobacco plants. The knockout of the endogenous *NtIAA4* gene (a *CsIAA4* homologous gene) in tobacco plants alleviated growth inhibition in *CsMYB4a* transgenic tobacco plants. Furthermore, protein–protein interaction experiments proved that domain II of CsIAA4 is the key motif for the interaction between CsIAA4 and CsMYB4a and that the degradation of CsIAA4 is prevented when CsMYB4a interacts with CsIAA4. In summary, our results suggest that CsMYB4a is a multifunctional transcription factor that regulates the auxin signaling pathway, phenylpropane and lignin pathways. This study provides new insights into the multiple functions of R2R3-MYB4 subgroup members as a group of well-known negative regulatory transcription factors.

**One-sentence summary:** CsMYB4a act as multifunctional transcription factor that regulates the auxin signaling pathway, phenylpropane and lignin pathways.

## Introduction

R2R3-MYB transcription factors belonging to the fourth subgroup (MYB4) are negative regulatory transcription factors of the phenylpropane and lignin pathways (Hemm et al., 2001; Ma and Constabel, 2019). Improving the taste of pasture by reducing the lignin content can increase the economic benefits of pasture. However, studies have found that these transcription factors that negatively regulate lignin synthesis also inhibit plant growth, thereby affecting their application potential in improving crop economic value (Jin et al., 2000; Li and Zachgo, 2013; Li et al., 2017). For example, to improve the quality of germplasm resources of lignocellulosic raw materials, PvMYB4 was overexpressed to reduce the lignin content of plants; however, plants with such overexpression exhibited traits of dwarfism and root growth inhibition (Shen et al., 2012).

A practical problem is how to achieve MYB4 to reduce the lignin content without reducing plant growth. MYB4 overexpression inhibits the transcriptional expression of phenylpropane pathway genes to reduce the lignin content but also causes growth inhibition, which is called lignin modification-induced dwarfism (LMID) (Panda et al., 2020). In a previous study, to promote the growth of MYB4 transgenic plants, its promoter was replaced, so that PvMYB4 was only overexpressed in green tissues, but not in the roots; this does not affect the growth of the above-ground parts of plants, while reducing their lignin content (Liu et al., 2018).

Recent experiments have found that the action sites of the fourth subgroup of R2R3-MYB transcription factors are both the phenylpropane and lignin pathways. CsMYB4a is belongs to the fourth subgroup of R2R3-MYB transcription factors and is derived from tea plants (*Camellia sinensis*). Its amino acid sequence is 60.44% identical to that of AtMYB4 (AEE86955). Transgenic tobacco plants with *CsMYB4a* overexpression exhibit abnormal growth compared with wild-type plants, with the symptoms of dwarfing, small leaves, and shrunken leaves (Li et al., 2017). *Cs*MYB4a negatively regulates not only the phenylpropanoid pathway but also the shikimate pathway in genetically modified tobacco plants. Some studies have shown that the synthesis and signaling pathways for the hormone ABA are the action sites of R2R3-MYB transcription factors. AcoMYB4 inhibits the transcriptional expression of genes involved in ABA synthesis and ABA signaling pathways, including *AcoABA1* and *AcoABI5* genes (Kim et al., 2015; Chen et al., 2020). A recent report is about the GROWTH INHIBITION RELIEVED 1 (GIR1) gene, its cloned protein is involved in the nuclear transfer of MYB4, and its destruction alleviates the growth inhibition in the Arabidopsis lignin mutant ref8, but did not restore the expression of phenylpropane pathway genes in the mutant(Panda et al., 2020). These results suggest that the partial role of MYB4 in the growth inhibition in genetically altered plants may be independent of the transcriptional regulation of phenylpropane biosynthesis genes.

In this study, we established a yeast two-hybrid library using CsMYB4a as the bait protein, and from the library, we discovered some proteins related to the auxin (AUX) signal transduction pathway, such as the regulatory factor AUX/indole-3-acetic acid (IAA) (**Table S1**). In addition to its role in phenylpropane and shikimic acid pathways, CsMYB4 may directly regulate the auxin signaling pathway, thereby affecting plant growth and development.

The AUX/IAA family of transcription factors comprises negative regulators of the auxin signal transduction pathway (Ulmasov et al., 1997; Leyser, 2018; Luo et al., 2018). The function of many AUX/IAAs has been studied and reported in recent years. *AtIAA3* inhibits the expression of *PIN* genes by interacting with *AtARF7* and *AtARF19*, affects the distribution of auxin in the root system of *Arabidopsis*, and inhibits the formation of lateral root primordia, leading to reduced lateral roots (Weiste et al., 2017; Orosa-Puente et al., 2018; Kumar Meena et al., 2019). The mutant of *AtIAA16* showed resistance to ABA during seed germination, and the plants showed insensitivity to auxin (Rinaldi et al., 2012). *OsIAA4-*overexpressing plants showed stunted and more tillering angles (Song and Xu, 2013). A knockdown mutant of *OsIAA6* showed abnormal tiller outgrowth (Jung et al., 2015). The expression of *OsPIN1b* and *OsPIN10a* was reduced in *Osiaa11*, and the development of lateral roots was inhibited (Zhu et al., 2012).

In this study, we speculate that CsMYB4a interacts with CsIAA4, which enhances the stability of the CsIAA4 protein and inhibits auxin signal transduction, which may have led to the repressed growth of plants. This regulation effect of CsMYB4a is independent of its regulation of the lignin synthesis pathway. First, we verified the interaction of CsIAA4 and CsMYB4a through protein–protein interaction experiments and found that domain II of CsIAA4 is the key site for the interaction. The degradation of the CsIAA4 protein was prevented under *CsIAA4* and *CsMYB4a* co-expression or their co-incubation *in vitro* and in tobacco plants. Our results reveal a new mechanism that MYB4 regulates auxin signaling transduction by directly binding to CsIAA4, preventing the CsTIR1-ubiquitin complex from ubiquitinating AUX/IAA, thereby inhibiting plant root growth. The results suggest that R2R3-MYB4 transcription factors are multifunctional and regulate the phenylpropane pathway and auxin signaling transduction.

## Results

### Tobacco plants overexpressing *CsMYB4a* exhibit abnormal responses to exogenous α-NAA

In this study, we first analyzed the expression differences of related genes in both auxin signaling pathway and phenylpropane pathway between transgenic tobacco seedlings and control seedlings (plants overexpressing empty vectors) at different developmental stages using transcriptome sequencing technology. Compared with control plants, the expression levels of some key genes in these two pathways showed a downward trend in transgenic plants, but the down-regulation of auxin signaling pathway genes occurs earlier than those of phenylpropane pathway genes. (**Fig. 1A, B**). The result suggests that CsMYB4a has a negative regulatory effect on plant growth that does not rely on the phenylpropane pathway. This means that *CsMYB4a* may directly regulate the auxin signaling pathway in addition to regulating the phenylpropane pathway and shikimic acid pathway, thereby affecting plant growth and development.

**Figure. 1.**
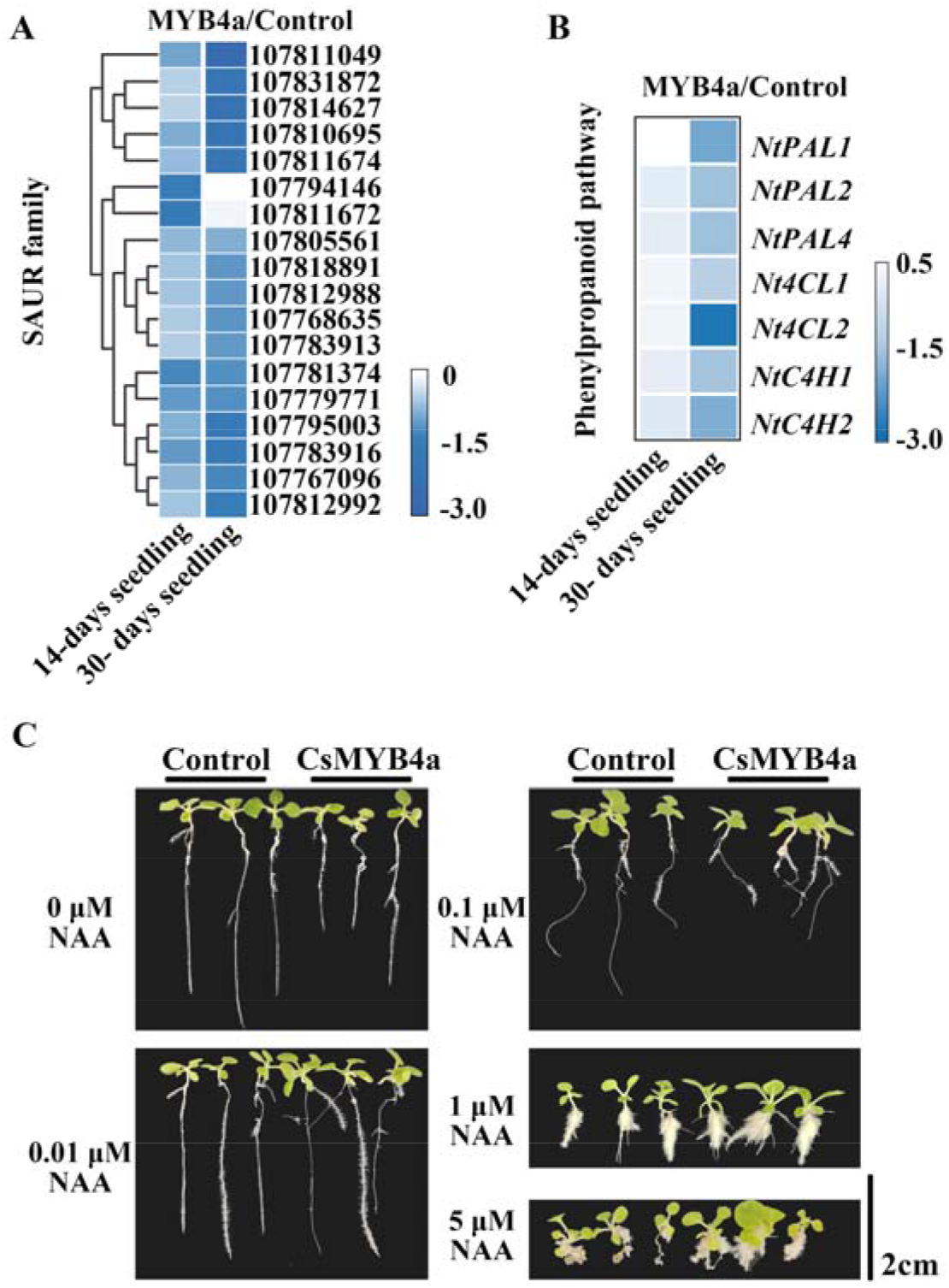
Effect of *CsMYB4a* gene on auxin signaling pathway in transgenic tobacco plants. A, the heatmap showing the expression differences of *Small auxin up RNAs* (*NtSAURs*) between transgenic tobacco seedlings and control seedlings at different developmental stages. The legend denotes log_2_ (fold change CsMYB4a/control). B, the heatmap showing the gene expression differences of the phenylpropanoid pathway between transgenic tobacco seedlings and control seedlings at different developmental stages. The legend denotes log_2_ (fold change CsMYB4a/control). C, morphological changes of control seedlings and transgenic seedlings after NAA treatment for 20 days. Scale bar = 2.0 cm.

Then, CsMYB4a transgenic plants (CsMYB4a) were exposed to a series of synthetic auxin 1-naphthaleneacetic acid (α-NAA) solvents at low to high concentrations, and their morphological changes were observed (**Fig. 1C**). With an increase in the hormone concentration from 0.01 to 5 μM, the root elongation of transgenic and control plants was suppressed, the roots gradually became shorter, and the number of adventitious roots increased gradually. Compared with control plants, the number of adventitious roots and the leaf area of transgenic plants increased significantly. This phenomenon was more significant under a high concentration of α-NAA.

### CsIAA4 is one of the target proteins regulated by CsMYB4

Yeast two-hybrid assay was performed to screen the target genes of CsMYB4 regulating the auxin signal pathway. Using *Cs*MYB4 as the bait protein, an AUX/IAA protein was screened out from the tea cDNA library. The unrooted PHYLIP phylogenetic tree showed the AUX/IAA family in tea plants contain 24 members, and this AUX/IAA protein was clustered on the same branch with *AtIAA1–4* proteins (**Fig. S1**). Based on its highest amino acid identity with *AtIAA4*, it was named *Cs*IAA4 (NCBI ID: ARQ20702.1). The expression data of AUX/IAA family in tea plants which extract from Tea Plant Information Archive (TPIA, <u>tpdb.shengxin.ren</u>) was showed in **Fig. S2**.

To explore the potential physiological significance of *CsIAA4* and *CsMYB4a* in the regulation in tea plants, the co-expression characteristics of the two genes in tea plants were studied using qRT-PCR experiments. The data revealed that *CsIAA4* and *CsMYB4a* were co-expressed in the different tissues and organs of tea plants; the expression data revealed that *CsIAA4* and *CsMYB4a* were highly expressed in mature flowers, petals, and primary roots (**Fig. S3A, B**). We performed a subcellular localization assay in tobacco leaves. The results showed that both *CsIAA4* and *CsMYB4a* were located in the nucleus of epidermal cells **(Fig. S3C)**.

*CsIAA4* was overexpressed in tobacco plants through the leaf disc method. The observed morphological features showed that the *CsIAA4* gene inhibited the growth and development of lateral roots and leaves of transgenic plants (**Fig. 2B**) and inhibited the seed germination of transgenic plants (**Fig. 2C**). However, the lignin content of genetically modified plants was not affected (**Fig. 2D**). This means that the overexpression of the *CsIAA4* gene does not disrupt the lignin synthesis pathway in transgenic plants.

**Figure. 2.**
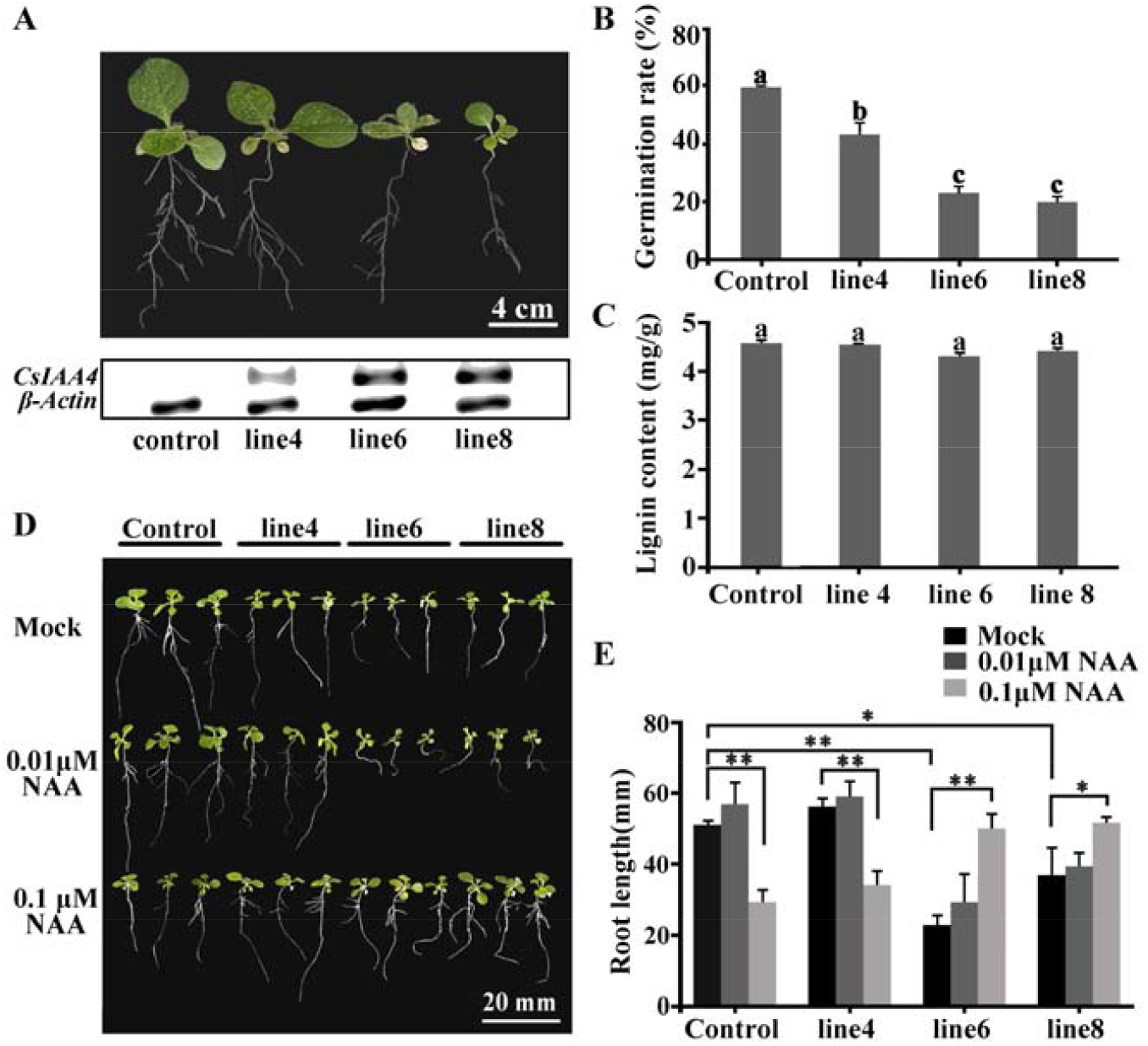
Effect of *CsIAA4* gene on the growth and response of transgenic tobacco plants to NAA treatment. A, morphological characteristics of a transgenic tobacco seedling (15 days old) overexpressing *CsIAA4*. Semi-qPCR showing the expression level of *CsIAA4* in a tobacco seedling. B and C, statistical data of plant growth and lignin content of a transgenic tobacco seedling (60 days old). Data are expressed as mean ± SD (n = 3), P < 0.05(Student’s *t* test). D, morphological changes of control and transgenic seedlings after NAA treatments for 20 days. Scale bar = 2.0 cm. E, statistical data of root elongation of transgenic plants in response to NAA treatment. Data are expressed as mean ± SD (n = 20), (*) P < 0.05, (**) P < 0.01(Student’s *t* test).

To test whether *CsIAA4* changes the response of plants to auxin, transgenic plants were exposed to exogenous α-NAA treatment at 0.01 and 0.1 μM (**Fig. 2E, F**). Morphological characteristics and statistical data showed that treatment with a low concentration of NAA promoted the root elongation of control plants, but the root elongation was inhibited under treatment with a high concentration of NAA. In comparison, the root elongation of transgenic plants with the high expression of the target gene (line6 and line8) did not change significantly under 0.01-μmol L^−1^ NAA treatment, but root elongation was significantly promoted under 0.1-μmol L^−1^ NAA treatment. NAA treatment changed the gravitational properties of the roots of the transgenic plants (line6 and line8) and the growth of the above-ground leaves. These results indicate that *CsIAA4* is a growth-inhibiting transcription factor, and some symptoms in response to exogenous α-NAA treatment of transgenic plants overexpressing *Cs*MYB4 and *Cs*IAA4 are consistent.

### CsMYB4a represses tobacco growth by regulating the auxin signaling pathway

To further investigate the interrelationship between *IAA4* and *CsMYB4a* on plant growth, we knocked out *NtIAA4* (Gene ID: 107830067), a homolog of *CsIAA4* (**Fig. S5A, B**), in control plants and *CsMYB4a*-line1 plants by using the CRISPR-Cas9 method. These mutants were named *ntiaa4* and *CsMYB4a*-*ntiaa4*, respectively. DNA sequencing results showed that some bases of the target gene were successfully knocked out in the two mutant plants (**Fig. 3A**). Morphological features showed that just knocking out *NtIAA4* in wild-type G28 plants did not promote plant growth (**Fig. 3B**), and statistical data also showed similar results (**Fig. 3C**). The growth of *ntiaa4* transgenic mutant was not promoted, suggesting that other AUX/IAAs with redundant functions may exist in tobacco plants.

**Figure. 3.**
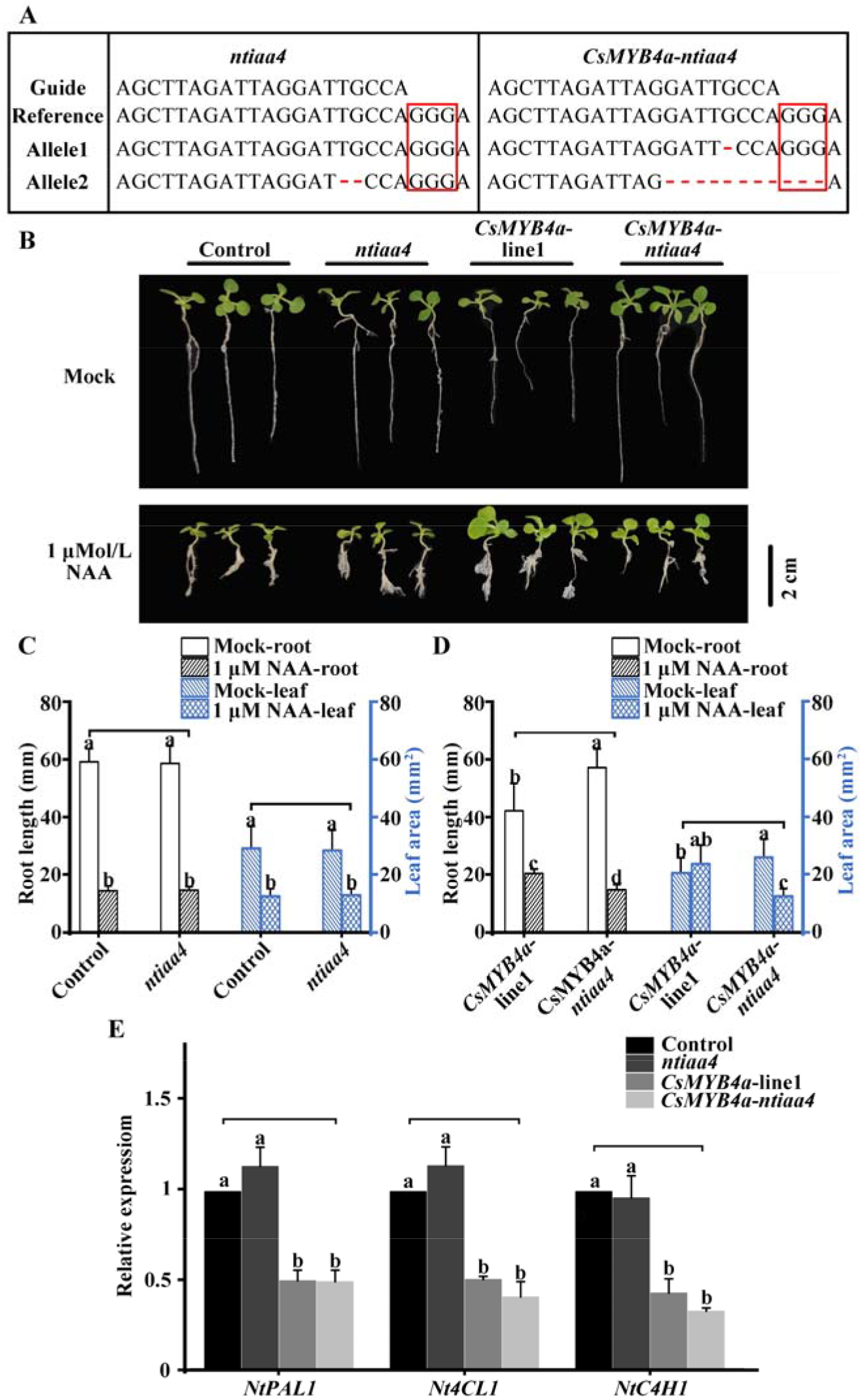
*NtIAA4* gene knockout experiment based on CRISPR-Cas9-mediated genome targeting system. A, the decoding of *NtIAA4* knock out plant superimposed sequencing chromatograms. B, morphological changes of control seedlings and *ntiaa4, MYB4a*-*ntiaa4* transgenic seedlings after NAA treatment for 20 days. Scale bar = 2.0 cm. C and D, Statistical data of root elongation and leaf area of transgenic plants response to NAA treatments. Data are expressed as mean ± SD (n = 20), P < 0.05(Student’s *t* test). E, Relative expression of the genes of the phenylpropane pathway in control seedlings and *ntiaa4, MYB4a*-*ntiaa4* transgenic seedlings. Data are expressed as mean ± SD (n = 3), P < 0.05(Student’s *t* test).

Compared with plants overexpressing *CsMYB4a*, the root elongation and leaf growth of *CsMYB4a*-*ntiaa4* plants with *NtIAA4* knocked out were significantly promoted **(Fig. 3B, C**), suggesting that *NtIAA4* knockout alleviated growth inhibition induced by MYB4. Moreover, the increase in the number of adventitious roots and the leaf area of *CsMYB4a-line1* plants promoted by the high concentration of NAA was not observed in *CsMYB4a*-*ntiaa4* plants.

To determine whether the lignin biosynthesis of *CsMYB4a*-*ntiaa4* transgenic plants is affected, we detected the expression of genes involved in the phenylpropane pathway. The result showed that the expression of *PAL1, 4CL1*, and *C4H1* of *CsMYB4a-ntiaa4* transgenic plants was similar to that of *CsMYB4a* transgenic plants MYB4a-line1 (**Fig. 3D**). The result indicated that the root elongation and leaf growth of *CsMYB4a*-*ntiaa4* plants are irrelevant to the lignin content.

The aforementioned results indicate that NtIAA4 is likely to be the target protein of CsMYB4a in transgenic tobacco plants.

### CsMYB4a interacts with the domain II of CsIAA4

To examine whether CsMYB4a interacts with CsIAA4, the pull-down assay, luciferase reporter (LUC) assay, and bimolecular fluorescence complementation (BIFC) assay were performed. *Cs*IAA4 contains four domains, as shown in motif locations of CsIAA4 in **Fig. S1B**. Domain II is involved in the regulation of ubiquitination, and it is suspected to be the site of interaction with CsMYB4a.Therefore, a truncated amino acid sequence was designed, in which domain II of CsIAA4 was knocked out, and the mutant protein was named CsIAA4D2 (**Fig. 4A**). Then, we performed *in vitro* pull-down assays using MBP-tagged CsMYB4a as the prey protein and His-tagged CsIAA4 and His-tagged CsIAA4D2 as the bait proteins (**Fig. S6A-C**). As shown in **Fig. 4A**, CsIAA4 pulled down CsMYB4a, but CsIAA4D2 could not interact with CsMYB4a *in vitro*. This result shows that domain II of CsIAA4 is the key motif for the interaction between CsIAA4 and CsMYB4a. The LUC assay showed that co-expressed CsIAA4-nluc and cluc-CsMYB4a proteins reconstituted LUC activity, but CsIAA4D2 could not (**Fig. 4B**). The results of the BIFC assay showed that YFP signals were reconstituted when CsIAA4 and CsMYB4a proteins were co-expressed. However, CsIAA4D2 could not (**Fig. 4C**). These assays suggest that CsMYB4a interacts with CsIAA4 *in vivo*, that the interacting site is domain II of CsIAA4, and that CsMYB4a may regulate the degradation of CsIAA4.

**Figure. 4.**
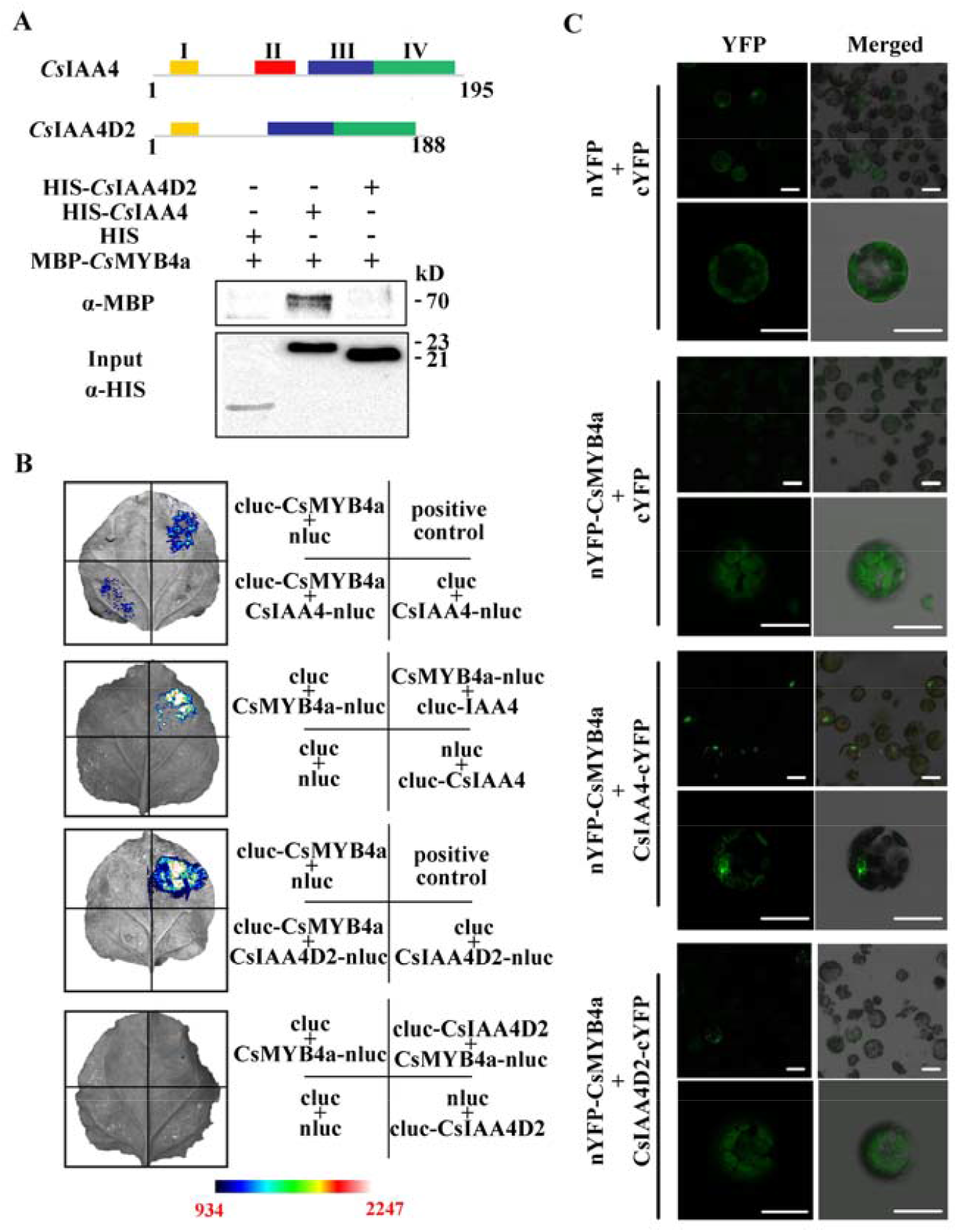
CsIAA4 interacts with CsMYB4a *in vitro* and *in vivo*. A, a diagram of the truncated experiment of the amino acid sequence of CsIAA4 and CsIAA4D2; the pull-down assays showing the interaction of CsIAA4 and CsIAA4D2 with CsMYB4a *in vitro*. HIS served as a control. Protein molecular weight of CsMYB4a-MBP is 70kD, protein molecular weight of CsIAA4 and CsIAA4D2 is 23kD and 21kD, respecti B, LUC assays showing the interactions of CsIAA4 with CsMYB4a in a tobacco plant. The scale bar denote wavelength of the *in vivo* image system was 934 to 2247. C, BIFC assays showing the interactions of *CsIAA4* with *CsMYB4a* in the protoplast of *Arabidopsis*. cYFP and nYFP empty vectors served as internal controls in each combination. Scale bar = 50 μm.

### CsMYB4a prevent the degradation of CsIAA4

To examine whether CsMYB4a prevents CsIAA4 degradation, several protein degradation experiments were conducted *in vitro* and *in vivo*. We first expressed IAA4-MYC in tobacco plants and extracted the crude protein from tobacco plants. The MBP-MYB4a protein was purified from recombinant *Escherichia coli* (BL21). Then the degradation rate of IAA4-MYC was investigated, the results showed that most of the IAA4-MYC protein was degraded within four hours, and the degradation of IAA4-MYC was inhibited when it was treated with MG132 (**Fig. 5A, B**). The tobacco crude protein containing recombinant IAA4-MYC and MBP or MBP-MYB4a was co-incubated with 0.1 μM NAA at 25°C for 4 h *in vitro*. The time the concentration of MBP-MYB4a in the two experimental groups was 0.5× and 1×, respectively. The MBP protein served as a control. The Western blot result showed that IAA4-MYC degradation was prevented after IAA4-MYC co-incubated with MBP-MYB4a. At a higher concentration of MBP-MYB4a, IAA4-MYC was degraded to a lower extent (**Fig. 5C**).

**Figure. 5.**
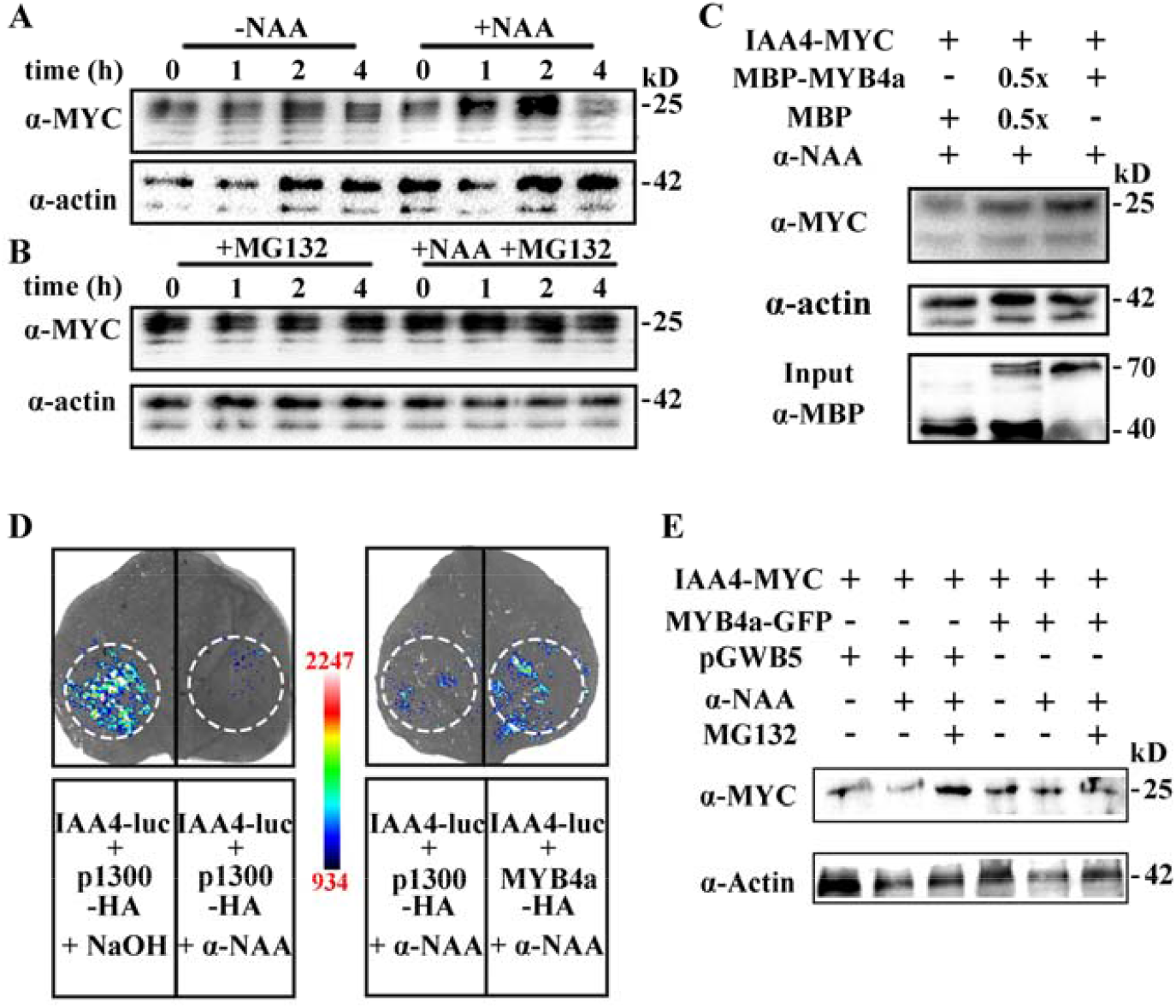
CsMYB4a prevents CsIAA4 degradation *in vitro* and *in vivo*. A, west-bolt showing the degradation of CsIAA4-MYC was caused by 0.1μM α-NAA *in vitro* at 4h, 25°C. CsMYC4a B, west-bolt showing MG132 repressed the degradation of CsIAA4-MYC under α-NAA treatment at 4h, 25°C. C, protein degradation assays showing that MBP-MYB4a prevents CsIAA4-MYC degradation in vitro when tobacco plants were treated with 0.1 μMol L−1 NAA. MBP protein served as a control. D and E, protein degradation assays showing that MYB4a prevents CsIAA4-MYC degradation *in vivo* when tobacco plants were treated with 0.1 μMol L^−1^ NAA. pGWB5 and p1300-HA empty vectors served as controls. The wavelength of the *in vivo* image system was 934 to 2247.

Using the transient expression system, CsIAA4-MYC and CsMYB4a-GFP were simultaneously expressed in tobacco plants, and these tobacco plants were then treated with exogenous NAA. The experiment of transforming CsIAA4-MYC and p1300-GFP were the control experiment. The Western blot results showed that exogenous NAA treatment promoted the degradation of the CsIAA4 protein when it was co-expressed with p1300-GFP in tobacco, and the degradation of the CsIAA4 protein expressed in plants was inhibited when it was co-expressed with CsMYB4a-GFP (**Fig. 5D**). A LUC detection experiment was also conducted to confirm that CsMYB4 can inhibit the degradation of CsIAA4 when CsIAA4-Luc and CsMYB4a-GFP were co-expressed in tobacco plants. Luc activity in plants expressing CsIAA4-Luc alone was reduced compared with that in plants co-expressing the two genes after α-NAA treatment (**Fig. 5E**). All the vector used for protein degradation were shown in **Fig. S6**.

In summary, the results showed that CsMYB4a interacts with CsIAA4 to prevent CsIAA4 degradation.

## Discussion

### CsMYB4a is a multifunctional transcription factor

MYB4 acts as an inhibitory transcription factor of the phenylpropane pathway (Jin et al., 2000; Hemm et al., 2001; Ma and Constabel, 2019) to not only inhibit lignin synthesis in MYB4-overexpressing tobacco plants but also inhibit the formation of lateral roots, delay plant growth (Shen et al., 2012), reduce sensitivity to auxin, and cause the development of long and narrow cotyledons, shrinkage and white spots leaves (Li et al., 2017). It is mostly believed that the growth inhibition caused by MYB4 overexpression is due to the reduction of lignin (Geng et al., 2020). MYB4 can inhibit the expression of PAL, C4H, COMT, and other genes in the phenylpropane metabolism pathway (Cavallini et al., 2015; Li et al., 2017), and the downregulation of these genes reduces the lignin content while causing growth defects (Huang et al., 2010; Saluja et al., 2021).But, CsMYB4a transgenic tobacco transcriptome data suggest that MYB4a regulates the auxin signal transduction pathway earlier than the phenylpropane pathway (**Fig. 1A, B**).

auxin is an important plant hormone that can regulate the growth and development of plants as well as the occurrence of lateral roots and adventitious roots, and it promotes cell differentiation (Lv et al., 2019). auxin promotes or inhibits plant growth through the TIR1-AUX/IAA-auxin response factor (ARF) signaling pathway in response to different environments (Leyser, 2018). The functions of AUX/IAA family genes are diverse, such as mediating embryo development (Hardtke et al., 2004), leaf shape and leaf color, root development (Swarup et al., 2008; Weiste et al., 2017), and photomorphogenesis (Yang et al., 2018). Regarding the underlying mechanism, AUX/IAAs use domains III and IV to interact with ARFs to inhibit the transcriptional expression of genes activated by ARFs (Tiwari et al., 2003; Guilfoyle and Hagen, 2007). In this study, CsIAA4 was screened from the yeast two-hybrid library of CsMYB4a and served as a negative regulator of the auxin signal transduction pathway. The expression of *CsIAA4* and *CsMYB4a* was consistently found in tissues and organs, and both proteins showed nuclear localization (**Fig. S3**). The results of the analysis of tissue and organ expression in Shuchazao plants showed that *CsIAA4* was highly expressed in petals, lateral roots, and old roots (**Fig. S3**); this expression pattern was positively correlated with the distribution of auxin in plants, indicating that CsIAA4 may be involved in auxin-induced lateral root occurrence and root growth. To examine the function of CsIAA4, we overexpressed *CsIAA4* in tobacco plants. We found that CsIAA4 acts as a suppressor of auxin signal transduction, and tobacco plants overexpressing this gene exhibited the characteristics of reduced lateral roots, slow growth, and reduced auxin sensitivity (**Fig. 2**). AtIAA1 and AtIAA3, which were located in the same branch of the phylogenetic tree as IAA4 in this study (**Fig. S1**), also regulate root development (Yang et al., 2004; Weiste et al., 2017; Orosa-Puente et al., 2018). In this study, the result of the exogenous auxin treatment assay showed that *CsIAA4* overexpression reduced the sensitivity of transgenic tobacco plants to auxin. High concentrations of auxin can recover the traits of reduced lateral roots and suppressed main root elongation in transgenic tobacco plants, suggesting that the transfer of *CsIAA4* into tobacco plants affects auxin signal transduction. Regarding transgenic tobacco plant traits, CsIAA4 and CsMYB4a promoted the traits of reduced lateral roots and delayed growth (**Fig. 1C** and **2A**). The growth inhibition of CsMYB4a transgenic tobacco plants was alleviated when NtIAA4 (homologous gene of CsIAA4) was knocked out in CsMYB4a tobacco plants (**Fig. 3**), indicating that CsMYB4a participates in the regulation of auxin signal transduction. By detecting the lignin content of transgenic tobacco plants, we found that CsIAA4 could not inhibit the synthesis of lignin (**Fig. 2C**). The findings indicate that CsMYB4a regulates the auxin signal transduction pathway; the underlying mechanism is independent of LMID induced by CsMYB4a (Shen et al., 2012).

### The model of MYB4a interacting with IAA4 to repress plant growth

At a low auxin concentration, AUX/IAAs recruit TOPLESS family proteins (TPLs), which are transcriptional co-repressors in plants and interact with ARFs to form heterodimers to inhibit the transcription activity of ARFs (Tiwari et al., 2003; Szemenyei et al., 2008; Leyser, 2018). At a certain concentration in plants, auxin promotes the ubiquitinated degradation of AUX/IAAs by promoting the targeted interaction of ubiquitin-ligase SCF^TIR1^ with AUX/IAAs (Gray et al., 2001). Domain II is a key domain that regulates the degradation of AUX/IAAs (Ramos et al., 2001; Lakehal et al., 2019; Kim et al., 2020), which leads to the elimination of the inhibition of the ARF transcriptional activity (Yang et al., 2004; Leyser, 2018; Luo et al., 2018). Some proteins prevent the degradation of AUX/IAAs by interacting with domain II of AUX/IAAs. For example, rice plants infected with the rice dwarf virus (RDV) showed dwarfing (Jin et al., 2016). The reason is that the RDV P2 protein interacted with domain II of *OsIAA10*; thus, *OsIAA10* cannot be ubiquitinated by *OsTIR1*. Similarly, in deep shade, phytochrome A (PHY A) competes with TIR1 by directly binding to the N-terminus of IAA3 and IAA17 to prevent the shade-induced degradation of IAA3 and IAA17, inhibiting the growth of hypocotyls (Yang et al., 2018). Cryptochrome 1 (CRY1) and phytochrome B (PHYB) also inhibit the growth of hypocotyls under blue and red light by interacting with Aux/IAA proteins (Xu et al., 2018; Mao et al., 2020). Due to the special structure of the CsIAA4 protein, the degradation of CsIAA4 can be mediated only when it is bound to domain II of CsIAA4. The protein interaction experiments in this study were conducted after removing domain II of CsIAA4. The result of the pull-down assay revealed that CsIAA4 interacts with CsMYB4a *in vitro*. The results of the Luc assay and BIFC assays suggest that CsIAA4 interacts with CsMYB4a *in vivo*, and the interaction site is the nucleus. It was also found that CsIAA4D2 does not interact with CsMYB4a (**Fig. 4**). The results confirmed that CsMYB4a interacts with CsIAA4 by combining with domain II of CsIAA4. To further verify whether the stability of CsIAA4 increases after CsMYB4a interacts with CsIAA4, protein degradation assays were performed (**Fig. 5**). The results showed that CsMYB4a prevents CsIAA4 degradation *in vitro* and *in vivo*.

Here, we propose a model (**Fig. 6**). When CsMYB4a expression increases during plant growth or when MYB4 transports MYB4 into the nucleus under drought or ABA regulation (Verslues et al., 2006; Zhao et al., 2007), CsMYB4a competes with CsTIR1 to interact with domain II of CsIAA4, which increases the stability of CsIAA4. When the auxin concentration increases, CsIAA4 cannot be degraded, which inhibits CsARFs from activating downstream auxin-responsive genes, and plant development is inhibited. CsIAA4 and CsMYB4a constitutive expression varies in organs and tissues of tea plants, except for leaves (**Fig. 2A**). The inhibition of the MYB4a-IAA4 complex does not always occur during plant growth; the repression may be eliminated when MYB4a is degraded by ubiquitination and phosphorylation. In *A. thaliana*, the expression level of *AtMIEL1* increases to degrade MYB30 for preventing programmed cell death during normal growth; the expression level of MIEL1 decreases, increasing the accumulation of MYB30 to resist the invasion of pathogens (Marino et al., 2019). MdMIEL1 negatively regulates the accumulation of anthocyanins by regulating the degradation of MdMYB1 protein (An et al., 2017). In *Populus*, the phosphorylation of PdLTF1 (an MYB transcription factor) by PdMPK6 is the key to regulate lignin accumulation in response to environmental stimuli (Gui et al., 2019). Secondary cell wall biosynthesis is regulated by MPK6 phosphorylation and the degradation of MYB46 in *Arabidopsis* (Im et al., 2021). Therefore, a MYB4a degradation pathway in plants may alleviate the inhibition of the CsMYB4a-CsIAA4 complex.

**Figure. 6.**
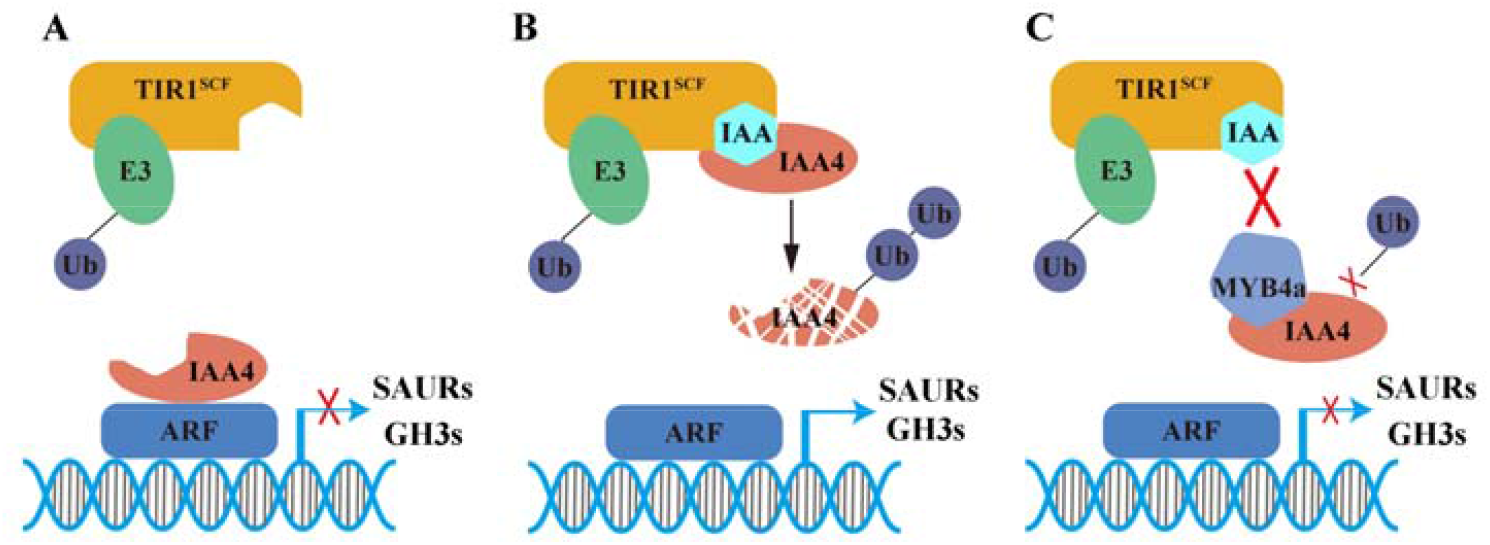
The model of the MYB4a regulation of CsIAA4 degradation. A, IAA4 interacts with ARFs to repress the auxin response gene. B, TIR1 interacts with IAA4 through auxin mediation and causes IAA4 degradation by ubiquitination; thus, the transcriptional activity of ARFs is alleviated. C, MYB4a interacts with the domain II of the CsIAA4 to compete with TIR1. The degradation of IAA4 is prevented when the expression of MYB4a increased.

## Materials and Methods

### Materials and growth conditions

All of the tea plants (*C. sinensis var. sinensis cv. Shuchazao*) and tobacco plants (*Nicotiana tabacum, common tobacco*) used in this study were grown in MS plates or soil at 25°C. CsMYB4a and CsIAA4 were cloned into a pCB2004 vector, which was transferred into tobacco plants through the *Agrobacterium*-mediated leaf disk transformation method. All primers used for vector construction in this study are shown in **Table S2**.

### Plant total RNA extract and qRT-PCR assay

A sample and an equal volume of polyvinylpyrrolidone (PVPP) were added in a mortar and ground to powder in liquid nitrogen, and 100 mg of powder was then transferred to a 2-mL RNase-free tube. Subsequently, 1 mL of Fruit-mate (Takara, catalog no.: 9192) was added, incubated at 37°C, centrifuged at 12,000 rpm for 5 min at 4°C. Subsequently, 400 μl of supernatant was transferred to a new tube, and 400 μl of RNAiso plus (Takara, catalog no.: 9108) was added for RNA isolation and centrifuged at 12,000 rpm for 5 min at 4°C. Subsequently, 160 μl of CHCl_3_ was added and centrifuged at 12,000 rpm for 15 min at 4°C. Thereafter, 400 μl of the supernatant was transferred to a 1.5-mL tube, and 400 μl of isopropanol was added, incubated on ice for 5 min, and centrifuged at 12,000 rpm for 5 min at 4°C; RNA was washed twice used with 1 mL of ethanol. A rotary steamer was used to remove ethanol, and 30 μl of RNase-free water was added to dissolve the total RNA. The qRT-PCR primers were designed based on the MIQE guidelines and are listed in **Table S2**. PCR programs were performed as described previously (Wang et al., 2018).

### CRISPR-Cas9

The vector used for CRISPR-Cas9 was pCBSG, CRISPR-GE was used to the guide-RNA design (Xie et al., 2017). Guide-RNA were cloned into the pCBSG by T4 DNA ligase. The recombination vectors were transferred into Agrobacterium (GV3101, cover pSoup-p19, weidi bio), and transferred into tobacco plants through the leaf disc method. The DNA for sequencing was extracted by DNA extract kit (Tsingke Biotechnology Co., Ltd, Beijing, China). The decoding of superimposed sequencing chromatograms by using DSDecode (http://skl.scau.edu.cn/dsdecode/) (Liu et al., 2015).

### Recombination protein purification and *in vitro* pull-down assay

*CsIAA4, CsIAA4D2*, and *CsIAA4D34* c pRSFDuet™ vector, and *Cs*MYB4a was cloned into the pMAL-c2X vector. The recombination vectors were transferred into *E. coli* (BL21) for protein expression. HIS-CsIAA4, HIS-CsIAA4D2, HIS-CsIAA4D34, and MBP-CsMYB4a were purified following the instructions of Novagen and NEB. *CsIAA4* and *CsIAA4D2* were cloned into the pRSFDuet™ vector, and CsMYB4a was cloned into the pMAL-c2X vector. The recombination vectors were transferred into BL21 (*E. coli*) and used for protein expression. HIS-CsIAA4 and HIS-CsIAA4D2 and MBP-CsMYB4a were purified following the instructions of Novagen and NEB. The HIS, His-CsIAA4 and HIS-CsIAA4D2 proteins served as bait and were added to the affinity chromatography column containing 2 mL of His-Tag nickel filler. The columns were incubated at 4°C for 3 h and washed three times with PBS buffer (0.2 M NaH_2_PO_4_, 0.2 M Na_2_HPO_4_, and 0.3 M NaCl, pH = 5.7). The MBP-CsMYB4 protein served as prey and was added to HIS, His-CsIAA4 and HIS-CsIAA4D2, and incubated overnight. Then, the fillers were washed three times with PBS buffer. Finally, the protein was eluted with 10 mL PBS elution buffer (0.2 M NaH_2_PO_4_, 0.2 M Na_2_HPO_4_, 0.3 M NaCl, and 1 mM imidazole, pH = 5.7), and the solution was added to a 10-kD protein ultrafiltration concentrator and centrifuged at 5500 rpm. The prey protein was analyzed through Western blotting with anti-HIS and anti-MBP (TransGen Biotech, Beijing, China).

### Dual-LUC assay

The vectors used for LUC assays were pC1300-cLuc and pC1300-nLuc, which contain fragments encoding the C- and N-terminal halves of Luc (cLuc and nLuc, respectively); *CsIAA4, CsIAA4D2*, and *CsMYB4a* were cloned in pC1300-cLuc and pC1300-nLuc vectors. The recombination vectors were then transferred into *Agrobacterium* (GV3101, cover pSoup-p19, weidi bio). These *Agrobacterium* cells were cultured at 28°C until the OD_600_ reached 0.6–0.8 and were centrifuged at 5500 rpm for 6 min in 50-mL tubes. To resuspend *Agrobacterium* cells, cell pellets were vortexed twice by using 10 mL of fresh infiltrate buffer (10 mM MgCl_2_, 10 mM MES, and 0.1 mM acetosyringone, pH = 5.6). The OD_600_ of the suspension was adjusted to 0.8–1.0, and CsIAA4-nLuc and CsIAA4D2-nLuc were then mixed with pC1300-cLuc as negative control and nLuc-CsMYB4a as the test group. nLuc-CsIAA4, nLuc-CsIAA4D2, and nLuc-CsIAA4D34 were mixed with pC1300-nLuc as negative control and CsMYB4a-cLuc as the experimental group, respectively. nLuc-CsCHI and CsDFR-cLuc served as the positive control. The positive control, two negative controls, and the experimental group were injected into tobacco leaves and occupied a quarter of the leaf, respectively. Tobacco plants were incubated in the dark for 48 h and then incubated under white light for 16 h. the leaves were cut off and spray D-luciferin, potassium salt on tobacco leaves and Luc fluorescence were detected using an *in vivo* imaging system.

### BIFC assay

The vectors used for BIFC assays were pBI221-cYFP and pBI221-nYFP, which contain fragments encoding the C- and N-terminal halves of YFP (cYFP and nYFP, respectively). The recombination vectors were transferred into *Agrobacterium* (GV3101, cover pSoup-p19, weidi bio). nYFP-CsIAA4, nYFP-CsIAA4D2, and nYFP-CsIAA4D34 were mixed with pBI221-cYFP as negative control and with CsMYB4a-cYFP as the test group. CsIAA4-cYFP, CsIAA4D2-cYFP, and CsIAA4D34-cYFP were mixed with pBI221-nYFP as negative control and with nYFP-CsMYB4a as the test group. *Arabidopsis* protoplast extraction and transfection were conducted following the protocol designed by Filip Mituła (Mituła et al., 2015). YFP signals were detected through confocal microscopy (Leica, DM2000).

### Protein degradation assays *in vitro* and *in vivo*

The vectors used for protein transient expression were p1300-MYC and p1300-HA (**Fig. S7**). For the *in vitro* assay, CsIAA4-MYC was transiently expressed in tobacco plants for 3 days; the leaves were then cut, and the same weight of PVPP was added to the leaves and ground in liquid nitrogen. Subsequently, 10 mL PBS (0.2 M NaH_2_PO_4_, 0.2 M Na_2_HPO_4_, 0.3 M NaCl, and 1 mM PMSF, pH = 5.7) was added to the powder, with additional grinding for 8 min. This PBS solution was transferred to a 50-mL centrifuge tube and centrifuged at 6500 rpm. The supernatant was transferred to a new centrifuge tube, and 85% (w/v) ammonium sulfate was added, shook on a decolorization shaker for 2 h, and then centrifuged at 6500 rpm for 10 min. The precipitate was dissolved in 1 mL of PBS and centrifuged at 6500 rpm for 10 min. IAA was added to the precipitate. MBP and MBP-MYB4a were purified using BL21. Subsequently, 20 μg of crude protein and 20 μg of MBP or 20 μg of MBP-MYB4a were co-incubated with 0.1 μM NAA at 25°C for 4 h, and the content of IAA4-MYC was detected through Western blotting.

For the *in vivo* assay, CsIAA4-MYC and CsMYB4a-GFP were co-expressed in tobacco plants for 3 days, and p1300-HA served as a control. Moreover, 0.1 μM NAA was sprayed on the leaves for 4 h before cutting. The crude protein was extracted, and the content of IAA4-MYC was detected through Western blotting.

### Protein degradation assays by Luc in tobacco

An additional 35S promoter was inserted before the Luc cassette of pGreenII 0800, and CsIAA4 was then cloned in the reconstructive vector pGreenII 0800 (**Fig. S7**) (Xu et al., 2018). The recombination vectors were transferred into *Agrobacterium* (GV3101, cover pSoup-p19, weidi bio). The method of tobacco transient expression was the same as that of the Luc assay.

## Acknowledgements

This work was financially supported by the Natural Science Foundation of China (32072621, 32002088, 31870676,) and the National Key Research and Development Program of China (2018YFD1000601).

